# Aberration correction for deformable mirror based remote focusing enables high-accuracy whole-cell super-resolution imaging

**DOI:** 10.1101/2023.12.01.568667

**Authors:** Wei Shi, Yingchuan He, Jianlin Wang, Lulu Zhou, Jianwei Chen, Liwei Zhou, Zeyu Xi, Zhen Wang, Ke Fang, Yiming Li

## Abstract

Single-molecule localization microscopy (SMLM) enables three-dimensional (3D) investigation of nanoscale structures in biological samples, offering unique insights into their organization. However, traditional 3D super-resolution microscopy using high numerical aperture (NA) objectives is limited by imaging depth of field (DOF), restricting their practical application to relatively thin biological samples. Here, we developed a unified solution for thick sample super-resolution imaging using a deformable mirror (DM) which was served for fast remote focusing, optimized point spread function (PSF) engineering and accurate aberration correction. By effectively correcting the system aberrations introduced during remote focusing and sample aberrations at different imaging depths, we achieved high-accuracy, large DOF imaging of the whole-cell organelles [i.e. nuclear pore complex (NPC), microtubules, and mitochondria] with a nearly uniform resolution of approximately 30 nm across the entire cellular volume.

## 1. Introduction

As a powerful tool for imaging biological samples, single-molecule localization based super-resolution microscopy allows for optical observation of structural details with unprecedented resolution while providing in-situ multi-targets imaging capability by multi-color labels[1, 2]. By engineering the PSF with different shapes along the axial direction, SMLM can easily achieve 3D super-resolution imaging capability. The most widely used 3D super-resolution imaging method for determining the axial position of the single-molecule involves introducing astigmatism into the detection path using a cylindrical lens[3]. However, this technique has limited DOF when a high-NA objective is used (∼1 µm). Considering that the typical thickness of a mammalian cell is 5-10 µm[4], this limitation prevents imaging of the entire cellular organelles that can extend over several microns throughout the whole cell[5].

To image samples thicker than the DOF of high-NA objective using SMLM[6-8], different implementation approaches have been developed, such as z-scan with thin sections imaging[4, 9-11], multiplane imaging[12, 13], and engineered PSF with large DOF imaging[14-17]. Z-scan imaging normally requires physically moving the sample or objective to scan at the axial positions of cells and then combining optical sections to obtain the complete cellular structure. It is therefore prone to stitching artefacts due to vibrations caused by stage movement during repeated scans of different sections. Multiplane detection allows for simultaneous imaging of different focal planes along the axial direction to achieve volumetric imaging. However, compared to single plane imaging, multiplane detection often suffers from lower signal-to-noise ratio (SNR) since the total number of photons is distributed across different imaging planes[13]. Thick sample imaging can also be achieved by engineered PSF with large DOF without the need to scan the sample. There are two main ways to generate these engineered PSFs. One is to fabricate a transmission phase mask with specialized pattern[15], placed in the Fourier plane of the microscope. Another way is to employ a programmable phase modulator, such as spatial light modulator and DM. Although both ways can be used to engineer PSFs with desired DOF, programmable phase modulators are often preferred due to their flexibility and ability to correct for both system and sample-induced aberrations. Especially with the development of DM based optimal PSF engineering, we were able to design an optimal PSF with predefined DOF without compromising the loss of photons[17]. However, there is normally a trade-off between the localization precision and DOF of the PSF optimized. PSFs with longer DOF normally spread in a larger area which leads to a reduction in SNR and loss of localization precision.

In recent years, adaptive optics (AO) techniques have been widely applied in optical microscopy[18]. In an approach known as remote focusing, fast 3D imaging of thick samples can be achieved by dynamically adjusting the focal plane without physically moving the objective lens or sample[19-23]. This process can be executed at rates of a few KHz using a DM or other variable optical elements[19, 24, 25], avoiding the need for physical movement of sample or objective. In SMLM, this method has demonstrated the capacity to record volumetric data of whole cells up to 10 µm while maintaining axial sample stabilization using a standard widefield microscope equipped with a DM[20]. However, system aberrations often arise when applying the defocus term to the vibrational device[22, 26]. Additionally, the inhomogeneous refractive indices of biological tissues can lead to the blurring and distortion of single-molecule emission patterns[11, 27]. All these issues can give rise to imaging artifacts and compromise the achievable resolution of SMLM. Hence, accurate correction and modeling of these aberrations are key to acquire high-accuracy super-resolution images across the entire volume.

Here, we employed DM to engineer PSFs with optimal 3D localization precision with different DOFs (DMO PSF). Furthermore, the DM was used to fast remote focus to longitudinally record axial stacks of whole cells across an extended range with high-accuracy, without the need to physically move the sample. Crucially, we accurately calibrated and corrected system aberrations introduced during refocusing at different imaging planes. Finally, we used in-situ single-molecules to estimate the aberrations at each focal plane through global fitting of the pupil function. The globally fitted aberrations were then fed to DM for sample-induced aberrations correction. In both silicone oil and oil objectives, we achieved consistent high-resolution 3D reconstructions at different depths and obtained a nearly uniform lateral Fourier ring correlation (FRC) resolution (∼34 nm for silicone oil objective and ∼30 nm for oil objective).

### 2. Remote focusing principle and optical setup

As shown in Fig. 1(a), we modified a standard 3D SMLM system by incorporating AO components. Remote focusing based on AO is enabled by a DM, which is positioned in conjugate with the objective pupil plane. The DM has emerged as the most prevalent phase modulator for fluorescence detection due to its minimal photon loss and rapid wavefront control capabilities. In this design, the DM serves three purposes: PSF engineering, aberration correction, and focal position shifting for different imaging depths. For precise wavefront control, we meticulously calibrated the experimental DM influence function of each actuator using a Twyman–Green interferometer, which measures the surface deformation of the DM[17, 28].

**Fig. 1.**
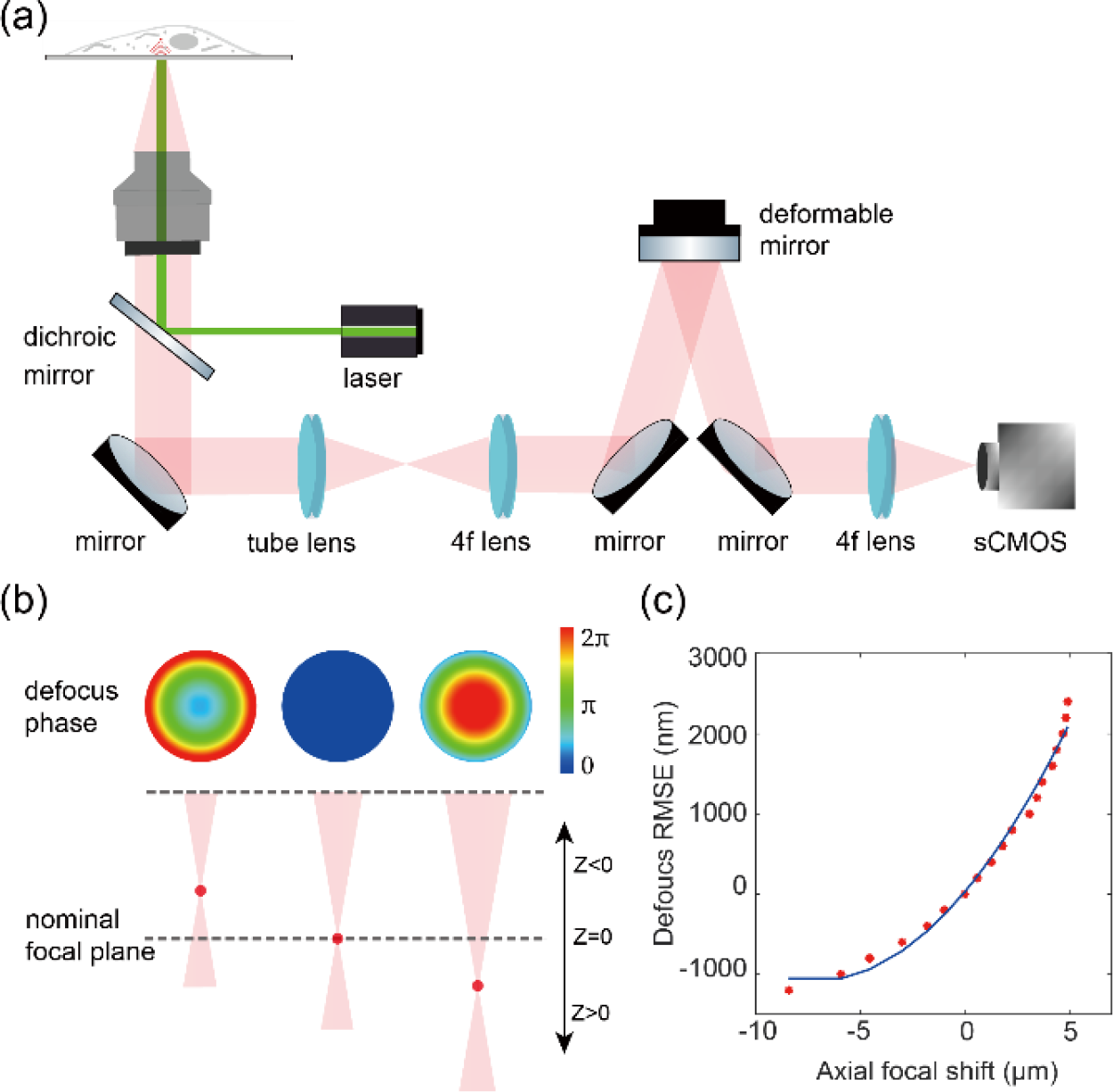
Schematic diagram illustrating the remote focusing principle. (a) Simplified schematic of a microscope configured for remote focusing using a DM in the imaging path. (b) The DM modulates the wavefront by controlling its defocus phase to shift different refocused focal planes around the nominal focal plane along the optical z-axis. (c) The defocus phase root mean square error (RMSE) for the optical wavefront is indicated when remotely focusing to different axial positions within the imaging optical system.

Remote focusing is a high-speed volumetric scanning technique that axially shifts the imaging plane by applying a defocus phase at the pupil of the optical system. This technique avoids the need to move the specimen or objective, thereby enabling accurate volumetric super-resolution imaging. We employed the defocus mode of the DM to refocus the focal plane of the imaging optical system. First, we defocused the beads on the coverslip using the objective’s z-piezo stage. Then, we compensated for this defocus by adjusting the DM’s defocus mode to refocus the beads. We calibrated the relationship between the amplitude of the applied defocus wavefront in DM and real axial shift of the imaging plane [Fig. 1(b)]. As shown in Fig. 1(c), the amplitude of the applied defocus wavefront doesn’t correlate linearly with the changes in the imaging plane. Especially for refocusing planes far away from the zero axial shift focal plane, the deviation is more pronounced. This is probably due to the limited actuators in the DM which cannot accurately modulate the ideal defocus wavefront. Therefore, it is very important to correct the residual aberrations at each focal plane after applying defocus wavefront to DM.

### 3. System aberration correction of remote focus using fluorescent beads

Astigmatism-based 3D SMLM imaging is the most widely used 3D super-resolution imaging method as it can be achieved by simply introducing a cylindrical lens into the imaging path. The DOF of the astigmatism PSF is approximately at the range of 1.2 µm around the focus[4]. Beyond this depth range, axial sample scanning[9] or optimized extended DOF PSF is necessary[29], as molecules outside the focus cannot be efficiently identified and localized. Furthermore, the 3D imaging capability of astigmatism is suboptimal as the localization precision decreases rapidly at the defocusing positions. In this study, we integrated our previously published DMO PSF[17] with remote focusing to achieve high-quality 3D super-resolution across the entire cell. To obtain the DMO PSF, we employed the influence function of the DM actuators as the solution space to optimize the pupil function of the engineered PSF by minimizing its 3D Cramér -Rao lower bound (CRLB). Subsequently, the DM was used to modulate both DMO PSF and astigmatism PSF [Fig. 2(a-d)]. Both experimental PSFs were built by axially scanning the beads on coverslip and interpolating the beads z stack with cubic spline[30]. The localization precision of both experimental PSFs was then compared. As shown in Fig. 2(e-f), the 2 µm DMO PSF exhibited a similar localization precision near the focus compared to the astigmatism PSF. However, the performance of astigmatism deteriorates significantly when the imaging plane is slightly away from the focus. In contrast, the DMO PSF maintains a relatively uniform resolution throughout the optimized axial range. The performance of experimental PSFs in Fig. 2(e, f) agrees well with the theoretical calculations[17].

**Fig. 2.**
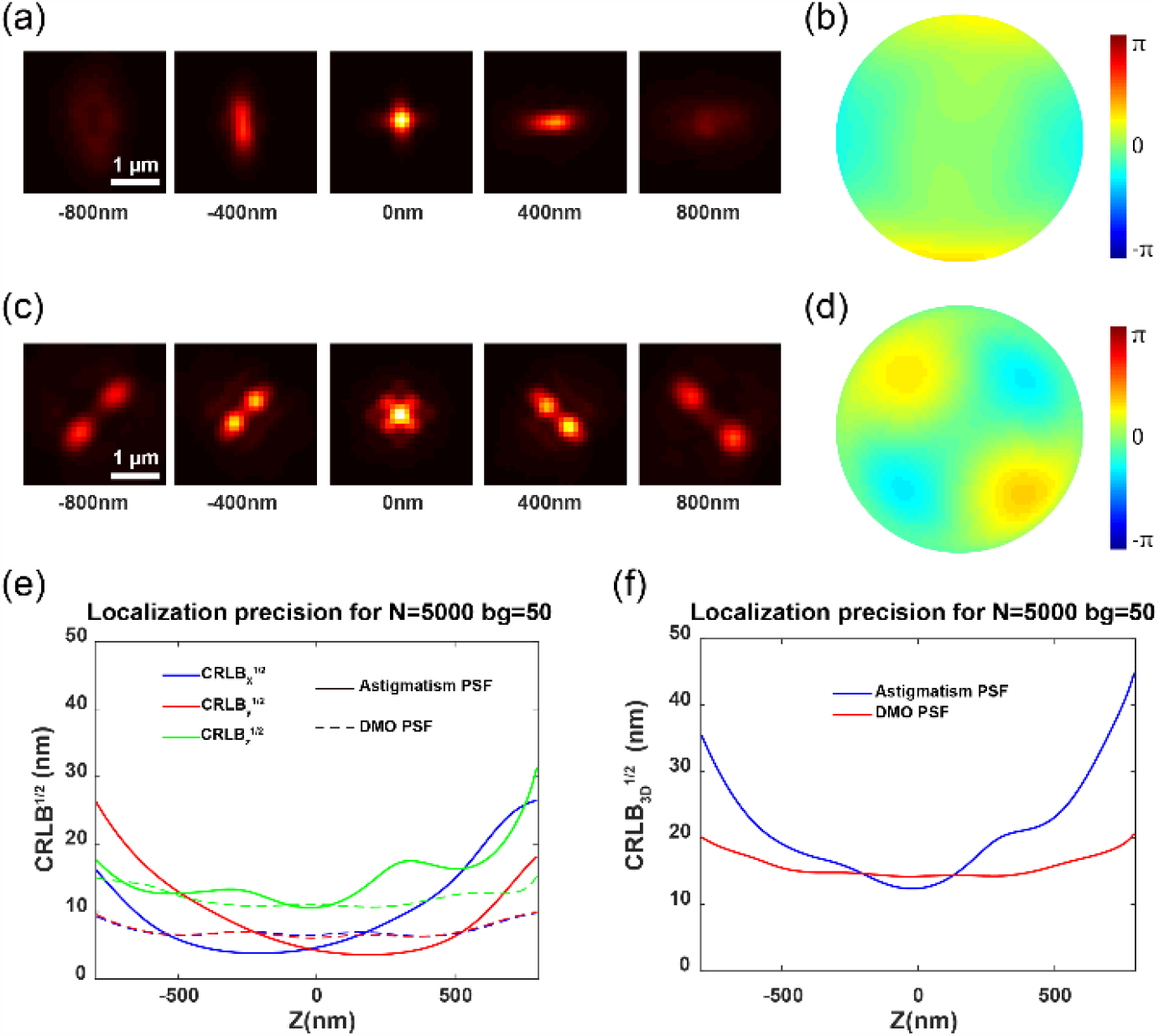
Comparison of the localization precision between experimental astigmatism PSF and DMO PSF. (a) Averaged astigmatism PSF of fluorescent beads on the coverslip, with PSFs at axial positions from -800 nm to 800 nm. (b) The pupil function corresponding to the astigmatism PSF of beads. (c) Averaged DMO PSF of fluorescent beads, identical to the range described in a. (d) Same as b for DMO PSF. (e) Comparisons of x, y and z localization precisions at different axial positions for both astigmatism PSF and DMO PSF. For the CRLB calculation, 5,000 photons and 50 background photons were used. (f) 3D CRLB of astigmatism PSF and DMO PSF.

We then applied the DM with the defocus wavefront in addition to the optimized pupil function to remotely scan the samples for volumetric super-resolution imaging. To investigate the PSF after adding the defocus wavefront, we imaged the beads on coverslip with DM-applied defocus wavefront and refocused the beads using the objective z stage. As shown in Fig. 3(a), the beads were modulated to the DMO PSF and defocus wavefront of -500 nm (4.62 rad, wavelength 680 nm) amplitude. The beads were then refocused by moving the objective 2 µm above the nominal focal plane. However, the refocused PSF showed significant difference compared to the one at zero-focus position [Fig. 3 (a) and Fig. 2 (c)]. We infer that this is probably due to the mismatch between the theoretical and experimental defocus mode, which leads to residual aberrations. To investigate the residual aberrations introduced beside defocus, we then acquired bead stacks in a range of ±1 µm relative t o the focal plane for calibration. We utilized a GPU-based vectorial PSF fitter to fit each bead stack by maximum likelihood estimation (MLE) to retrieve the Zernike-based aberration coefficients[16]. Furthermore, we employed globLoc[31] for the global fitting of images from the bead stacks, enabling high-accuracy estimation of aberrations.

**Fig. 3.**
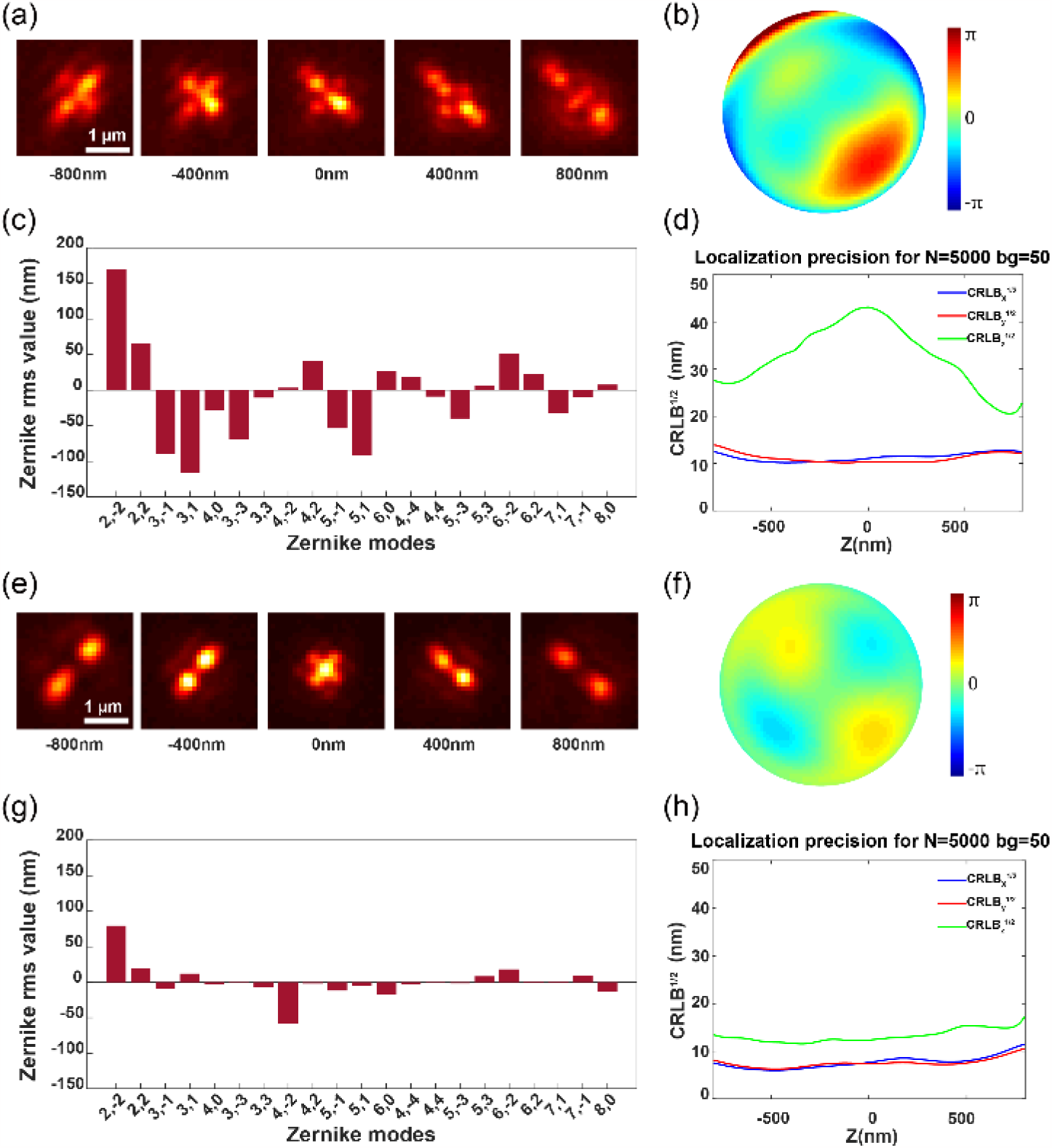
Correcting aberrations during refocusing with DMO PSF using a DM at various axial positions of the beads on the coverslip. (a) The beads were refocused using a DM to a position 2 µm above the nominal focal plane. (b) The pupil function for the refocused beads. (c) Fitted 21 Zernike coefficients for the beads stack. (d) CRLB for the experimental DMO PSF, with 5.000 photons and 50 background photons used in the CRLB calculation. (e-h) The PSF shape, pupil function, Zernike coefficients, and localization precision after aberration correction, respectively. The CRLB exhibits significant improvements, particularly in the z localization precision.

In this study, we retrieved the coefficients of all 21 tertiary Zernike polynomials from the average experimental bead stack [Fig. 3(c)]. Due to the extra aberrations introduced besides defocus, the pupil function of the refocused DMO PSF [Fig. 3(b)] is quite different from that at the zero focal plane [Fig. 2(d)]. The residual aberrations also led to the degradation of localization precision, particularly in the z-axis direction [Fig. 3(d)]. Therefore, we compensated for the 21 Zernike aberrations using the DM to counteract the additional aberrations introduced after refocusing, making the aberrations at the refocusing position closer to those of the beads on the coverslip [Fig. 3(e) and 3(f)]. As shown in Fig. 3(h), the resulting localization precisions are close to those of the PSF at zero focal plane [Fig. 2(e)].

We then demonstrated the effectiveness of the residual aberration correction after applying defocus wavefront using the nucleoporin Nup96 in U2OS cells, a structure often used as a quantitative reference [32]. Top nuclear envelope of the Nup96-SNAP-AF647 labeled NPCs were imaged. The imaging depth is about 4 µm above the coverslip. As shown in Fig. S2(a), the single-molecule images are very aberrated and the SNR is considerably low due to the additional aberrations introduced when applying the defocus wavefront to the DM. We then applied the calibrated residual aberration correction using the beads on the coverslip as described before. After aberration correction, the single-molecules are clearly visible, and the SNR of the image is improved significantly [Fig. S2(b)]. The number of localized single-molecule events increased sixfold, and the signal background ratio (SBR) of single-molecule events improved by nearly 2.5 times [Fig. S2(c)], with the localization and quantification analysis of single-molecule images implemented in SMAP[33]. The experimental astigmatic PSFs with and without residual aberration correction after applying defocus wavefront are shown in Fig. S3. We then applied these two PSFs to image biological samples. As shown in Fig S4, the double ring structure of NPCs can be clearly resolved after residual aberration correction [Fig. S4 (f)] while it was hardly resolved when only defocus wavefront was applied to DM [Fig. S4(c)].

### 4. Whole-cell 3d imaging with remote focusing using beads PSF model

We first demonstrated the whole-cell imaging capability of our approach using a silicone oil objective which we can match the refractive index of the sample medium with silicone oil[34, 35]. Whole-cell nucleus was imaged with 2 µm DMO PSF. 5 imaging planes spaced by 1 µm were recorded by remote focusing between different imaging planes using DM (Methods). The 5 optical section data were then stitched together by redundant cross-correlation to reconstruct the whole nucleus (Methods). As shown in Fig. 4, we were able to clearly reconstruct the ring structure in both top and bottom nuclear envelope with FRC resolution of 34 nm [Fig. 4(b)]. In the side-view of the nucleus, we were also able to resolve the double ring structure in the entire nucleus over a depth of ∼5 µm. Moreover, both the reconstructed diameter and the spacing between the upper and lower rings of the Nup96 structure agree with the reference structure [Fig. 4(h-i) and 4(j-k)]. We then imaged the nucleus with 6 µm DMO PSF (Fig. S5) for comparison and observed a reduced FRC resolution (∼ 50 nm, Fig. S6). It is also difficult to observe the double ring structure of Nup96 from the side-view images.

**Fig. 4.**
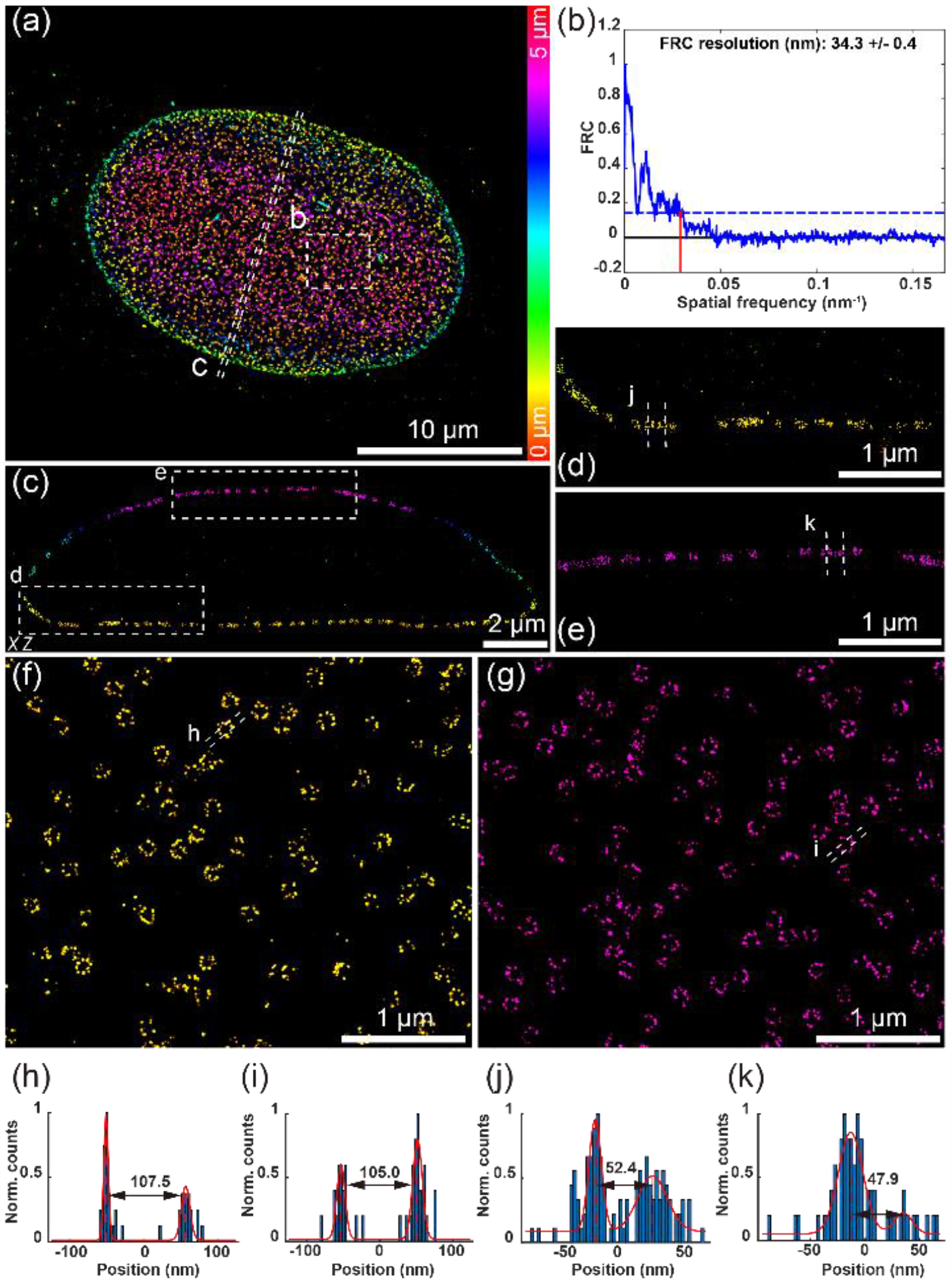
Whole-cell 3D super-resolution imaging of NPC. (a) Overview of the panoramic whole-cell 3D imaging of NPC using DMO PSF, merging 5 optical sections. (b) FRC analysis of the region boxed in a. (c) Side-view cross-section of the region denoted by the dashed line in a. (d-e) Magnified view of the area denoted by the box in c. (f) View of the bottom surface of the boxed area in a. (g) View of the top surface of the boxed area in a. (h-i) Intensity profile along the white dashed lines in f and g. (j-k) Intensity profile along the white dashed lines in d and e. The data were acquired from 8,000 frames per cycle over 20 cycles from 5 optical sections, with 120 mW laser power and 20 ms exposure time.

To further demonstrate that image quality is maintained throughout the entire cell, we imaged microtubules in COS-7 cells. The 3D reconstruction was assembled from 7 optical sections with step sizes of 1 µm, resulting in a whole-cell microtubule super-resolution image with a DOF of 8 µm [Fig. 5(a)]. We were able to clearly observe the microtubule structures throughout the whole cell, with the ability to resolve finer structural details of the microtubules as shown in both top and bottom layers of the cell [Fig. 5(b) and (c)]. Benefitting from the stitching method by redundant cross-correlation, no significant stitching artifacts were observed in the axial profile of the whole-cell microtubule volume, which is aligned of 7 optical sections [Fig. 5(d)].

**Fig. 5.**
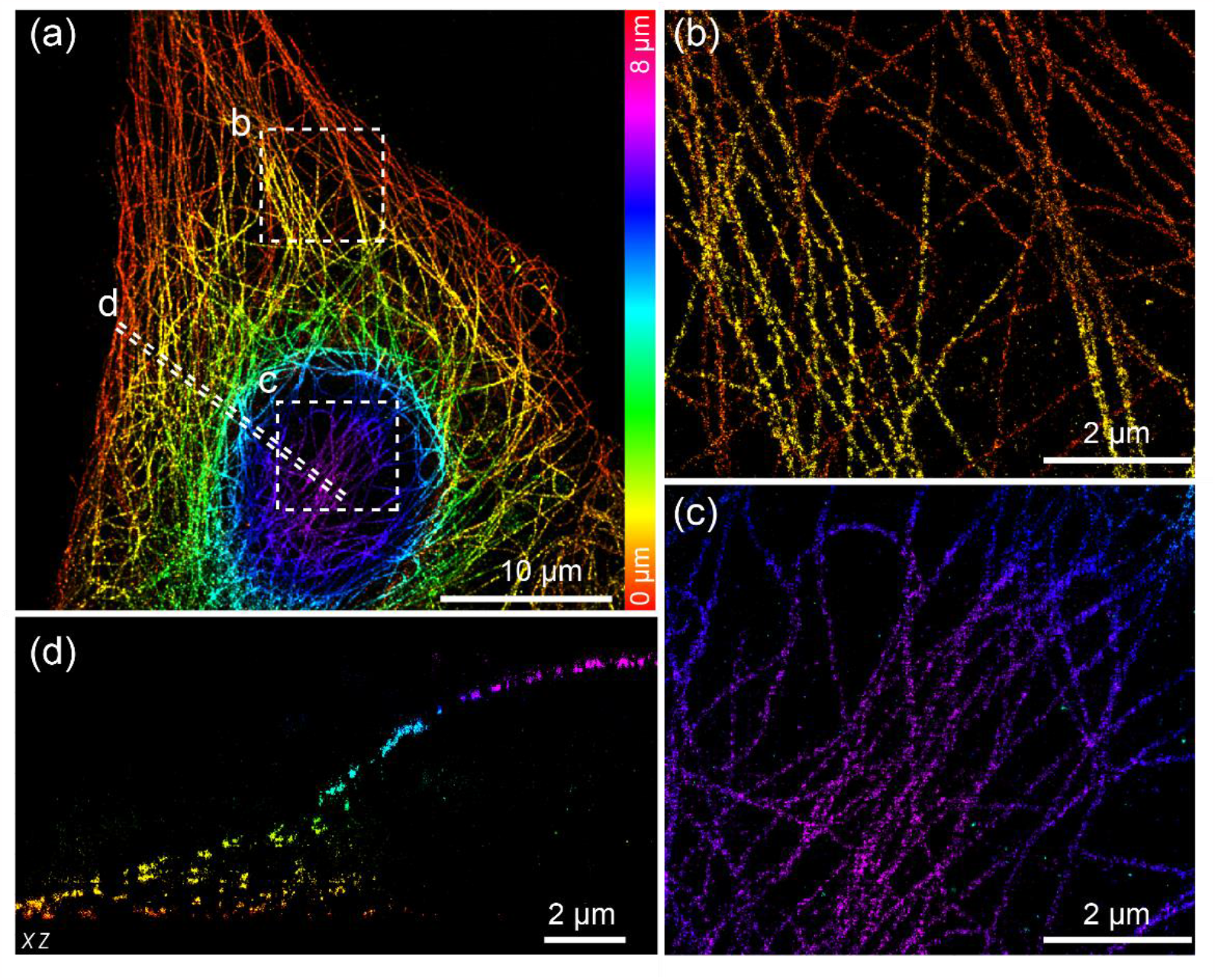
Whole-cell 3D super-resolution imaging of microtubules. (a) Overview of the panoramic whole-cell 3D imaging of microtubule using DMO PSF, merging 7 optical sections. (b) Zoomed bottom surface view of the boxed area denoted in a. (c) Zoomed top surface view of the boxed area denoted in a. (d) Side-view cross-section of the region denoted by the dashed line in a. The data were acquired from 7,000 frames per cycle over 40 cycles from 7 optical sections, with 200 mW laser power and 15 ms exposure time.

### 5. In-situ aberration correction of remote focus using blinking single-molecules

Bead-based PSF model normally works well only when the in-situ PSF model within the samples is close to the PSF model on coverslip. This is often valid when the refractive index of the sample medium is matched to the objective immersion medium. However, the refractive index of the sample medium is often not matched to the objective immersion medium. Especially for high-NA objective where oil is used as immersion medium, strong spherical aberrations will be introduced when the focal plane is away from the coverslip. These additional aberrations often result in reduced resolution if they are not corrected. To address this, we employed our recently developed in-situ PSF modeling method, uiPSF, to model the PSF directly from the single-molecule blinking events[36]. Subsequently, we carried out in-situ aberration correction for different imaging planes to further enhance the imaging resolution across the entire cell.

Here, we employed a 1.5 NA oil-immersion objective to image the whole cells. Similar to the silicone oil objective imaging, we first calibrated the residual aberrations at different focal planes after applying the defocus wavefront to the DM. As shown in Fig. 6 (a) and (b), an ideal DMO PSF could be obtained after applying the residual aberration correction. We then applied this PSF to image COS-7 cell at a depth of about 2.5 μm [Fig. 6(c)]. The single-molecule blinking data was used to estimate the in-situ PSF model. The pupil function was estimated by global fitting the single-molecule images via inverse modeling by uiPSF. As shown in Fig. 6(g), the fitted pupil function showed strong spherical and coma aberrations compared to the beads-based PSF on coverslip. This is probably due to the refractive index mismatch between the sample medium and oil immersion medium.

**Fig. 6.**
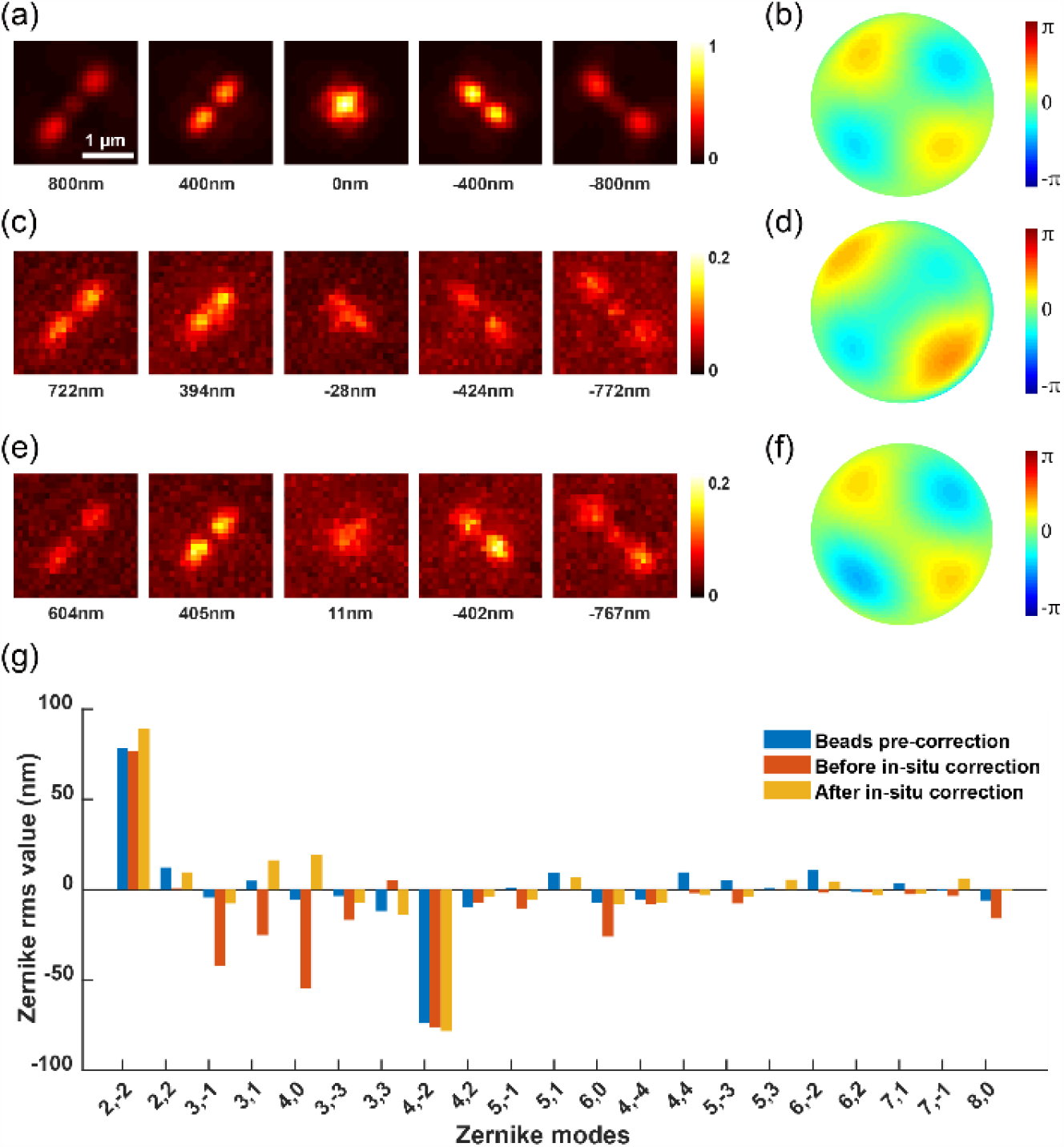
In-situ aberration correction using blinking single-molecules under remote focusing. (a) The beads on the coverslip were refocused using the DM to 1 µm above the nominal focal plane, with a pre-correction for aberrations applied to the imaging system. (b) Pupil function derived from the PSF of refocused beads. (c) The refocused PSF was obtained from single-molecule blinking data of immunofluorescence-labeled TOM20, recorded at a depth of 2.5 μm in COS-7 cells. (d) Pupil function calculated from the in-situ PSF. (e-f) The PSF shape and its corresponding pupil function after in-situ aberration correction. (g) The fitted 21 Zernike coefficients were obtained from the PSF of refocused beads after system aberration correction, the in-situ blinking single-molecule, and the in-situ PSF after aberration correction using blinking single-molecule.

Subsequently, we compensated for the aberrations by controlling the DM and then re-collected single-molecule data from the same location for in-situ PSF estimation. Through iterative corrections, various aberrations were effectively minimized [Fig. 6(g)]. The resulting single-molecule data displayed a 3D PSF shape [Fig. 6(e)] and pupil function [Fig. 6(f)] similar to those of the beads on the coverslip.

### 6. Whole-cell 3d imaging with remote focusing using in-situ PSF model

We then combined remote focusing with uiPSF based aberration correction using DM to image the whole-cell mitochondria using an oil objective. Here, we employed 2 µm DMO PSF and imaged the whole-cell mitochondria with 3 remote focusing imaging planes spaced by 1 µm. For each refocused position, in-situ PSF aberrations were corrected at depths of 0.5 µm, 1.5 µm, and 2.5 µm. Data from these 3 optical sections were acquired using rapid remote focusing cycles, controlled by the DM. The reconstructed 3D image was assembled from 3 optical sections, which were then aligned with adjacent optical sections, resulting in a whole-cell mitochondria super-resolution image with a DOF of 3 µm [Fig. 7(a)]. With our approach, the interconnected mitochondrial network was clearly resolved, and the x–z cross-sections revealed the membrane contour of mitochondria in the axial direction [Fig. 7(b)]. Thanks to the improvement in resolution through in-situ aberration correction across the whole cell, we achieved consistently high-resolution 3D reconstructions of organelles at different depths [Fig. 7(c-e)]. Resolution measurements were performed in regions proximate to both the lower [Fig. 7(f)] and upper [Fig. 7(g)] surfaces of the cell using the FRC method, and we obtained a nearly uniform lateral resolution of approximately 30 nm.

**Fig. 7.**
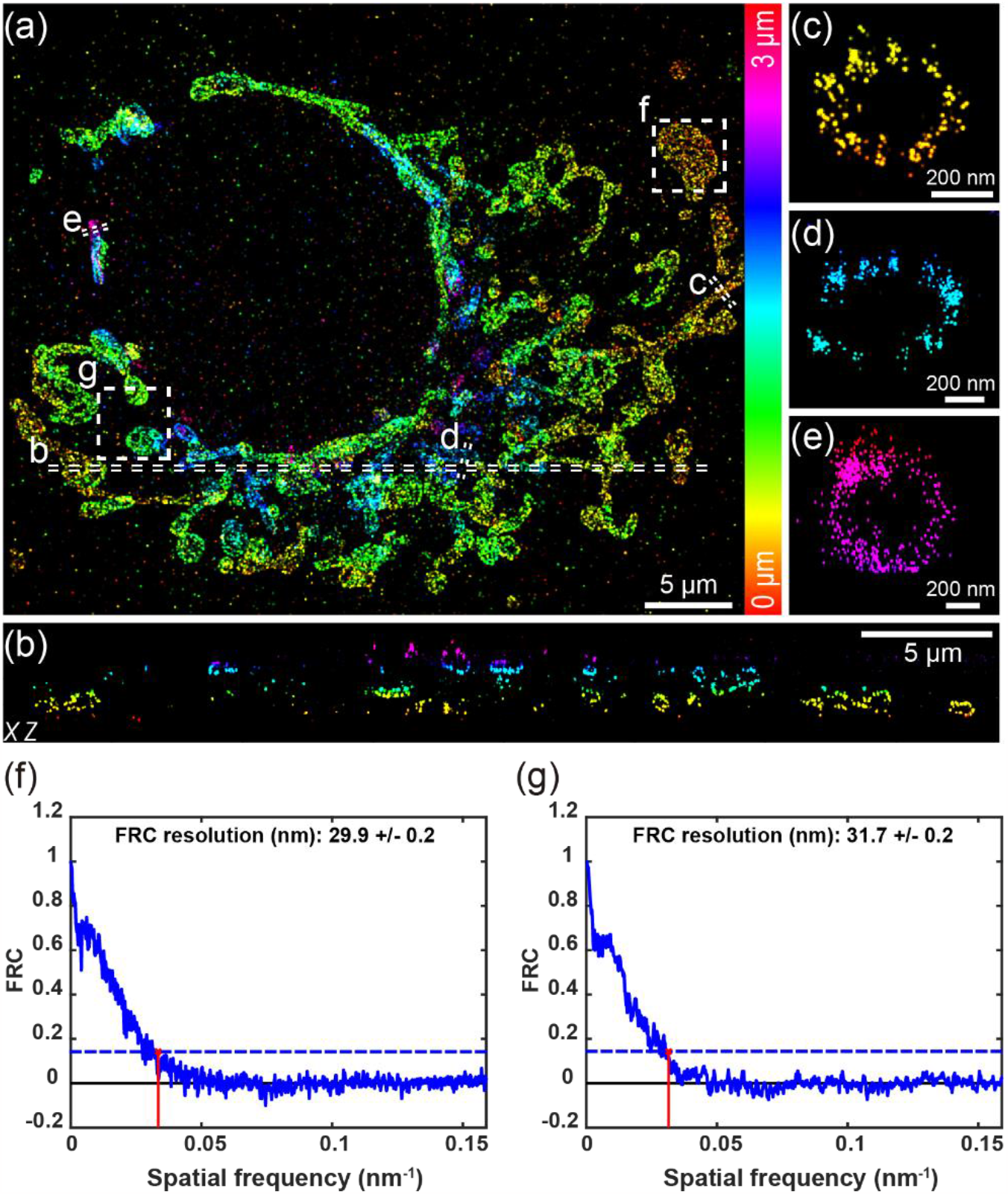
Whole-cell 3D super-resolution imaging of mitochondria. (a) Overview of the panoramic whole-cell 3D imaging of mitochondria using DMO PSF by merging 3 optical sections. (b) Side-view cross-section of the region indicated by the dashed line in a. (c-e) Zoomed-in side-view cross-sections from distinct depth areas, as indicated by the dashed lines in a. (f-g) FRC analysis of regions enclosed by the boxes in a, with f near the cell’s bottom surface and g near the top. The data were acquired from 3,000 frames per cycle over 70 cycles from 3 optical sections, with 200 mW laser power and 15 ms exposure time.

## 7. Discussion

In this work, we employed DM to perform super-resolution imaging of the whole cells by remote focusing. We systematically corrected the residual aberrations after applying the defocus wavefront and sample-induced aberrations with uiPSF. Furthermore, we utilized the DMO PSF, which could achieve optimal 3D resolution within a predefined DOF using DM. These resulted in a significant SNR improvement in single-molecule data post aberration correction. We demonstrated the performance of our approach by conducting whole-cell imaging of nucleoporinNup96, microtubules and mitochondria. Compared to the PSF directly optimized by the DM with a large DOF of 6 µm, which exhibited a resolution of approximately 50 nm, our reconstructed super-resolution images exhibited a marked improvement in resolution (∼30 nm). Moreover, we employed uiPSF to precisely estimate the sample induced aberrations using the in-situ blinking single-molecules when a high-NA oil objective was used.

In summary, the remote focusing enabled high-resolution whole-cell imaging, particularly in scenarios with high labeling density where large DOF PSF imaging could result in a strong background. With the precise in-situ PSF estimation, it can also be further extended to tissue imaging[37, 38]. Although we showed superior performance of remote focusing compared to the whole-cell imaging with large DOF PSF, it is worth noting that there is a balance between the number of focus planes imaged and the number of molecules acquired due to photobleaching. A combination of light sheet illumination with remote focusing could minimize the photobleaching while maintaining high-resolution imaging. We believe that our study will promote the use of adaptive optics in SMLM techniques, especially in biological applications that require both a large DOF and high-resolution imaging.

## 8. Methods

### A. Optical setups

In this work, we performed SMLM imaging at room temperature (24 °C) on a custom-built microscope (Fig. S1). We coupled the excitation light (iBEAM-SMART-405-S, 150mW, iBEAM-SMART-640-S-HP, 200mW, TOPTICA Photonics) into a single-mode fiber (P3-405BPM-FC-2, Thorlabs) using a dichroic mirror (DMLP 425, Thorlabs) and a fiber coupler (PAF2-A4A, Thorlabs). We adjusted the translation stage to vary the fiber output for different illumination angles. The illumination beam was filtered by a laser clean-up filter (ZET405/488/561/640xv2, Chroma) to remove excess wavelengths and stray light. We used a pair of lenses L1 (75 mm) and L2 (400 mm) with a slit (SP60, Owis) at AP1 for beam collimation and reshaping. The beam was then reflected by the main dichroic mirror (ZT405/488/561/640rpcxt-UF2, Chroma) before entering the objective for sample illumination. The emitted fluorescence was collected by a high-NA objective (NA 1.35, UPLSAPO 100 XS or NA 1.5, UPLSAPO 100 OHR, Olympus) and imaged by the tube lens TBL (TTL-180-A, Thorlabs) onto the AP2 confined by a slit (SP40, Owis). We utilized two bandpass filters (NF03-405/488/561/635E25, Semrock) and (FF01-676/37-25, Semrock) on the filter wheel to separate the emitted fluorescence from the excitation laser. We set up a 4f system (L3, 125 mm, L4, 75 mm) in the Fourier plane, with a deformable mirror (DM140A-35-P01, Boston Micromachines) for PSF engineering, aberration correction, and focal position shifting for different imaging depths. We could observe the objective’s back focal plane through lens L5 (75 mm) placed in front of the camera. Images were acquired using an sCMOS camera (Dhyana 400BSI V3, Tucsen) with a pixel size of 110 nm in the sample. Furthermore, we integrated a closed-loop focus lock control system into our optical setup. This system introduces the signal from a 785 nm laser (iBEAM-SMART-785-S, 125 mW, TOPTICA Photonics) reflected off the coverslip through a dichroic mirror (FF750-SDi02, Semrock) and detects it with a quadrant photodiode (SD197-23-21-041, Advanced Photonix Inc), which then provides feedback control to a z-piezo stage (P-726.1CD, Physik Instrumente) to maintain focus stability.

### B. Data acquisition

For calibrating the aberrations at different refocus positions, we acquired bead stack data on the coverslip after refocusing and modulated them to achieve a 2 µm DOF PSF using the DM. We acquired all bead stacks with an exposure time of 100 ms using a 642 nm laser at illumination intensities of approximately 10 mW. During sample aberration correction, we collected 10,000 to 30,000 frames of single-molecule data to the extract the in-situ PSF model and analyze Zernike coefficients based on the distribution density of the sample labeling structures at the refocus positions. To prevent fluorescence bleaching in cells during the iterative correction process, we performed preliminary corrections using separate cells. These pre-corrected Zernike coefficients were then applied during the actual imaging of other cells. When comparing the Zernike coefficients fitted from the actual imaging cells to the pre-corrected coefficients, only minor discrepancies were observed at the same in-situ depth (Fig. S7). While acquiring whole-cell data, we first determined the number of optical sections required based on the thickness of the cell under wide-field imaging, with each section spaced 1 µm apart. We then set the number of frames to be acquired for each section to 1,000-2,500 and the number of cycles to achieve a total of 20,000-70,000 frames at each section. The software control of the microscope was integrated into Micro-Manager with EMU[39], and we carried out data acquisition using Python scripts, leveraging the image acquisition interface of pycro-manager[40] to implement predefined DM acquisition modes. While using the beads PSF model, we imaged the sample in a refractive index matching buffer.

The refractive index of this buffer was matched to that of the silicone immersion oil (1.406), ensuring a constant refractive index axially. When employing the in-situ PSF model, we imaged the sample in a refractive index buffer that was not matched to the immersion oil (1.518). The sample was excited using a 642 nm laser with an intensity of 120-200 mW, and the 405 nm laser was adjusted according to the density of blinking single-molecule emitters. Typically, we acquired 100,000-300,000 frames per dataset with an exposure time of 15-20 ms.

### C. Data processing and analysis for multi-optical section

We have provided an open-source software for analyzing bead stacks and employed a GPU-based vectorial PSF fitter to fit them using MLE. The pupil function of average beads was decomposed into Zernike polynomials and calculated the coefficients of these polynomials. Additionally, we used an open-source deep learning-based algorithm to analyze the single-molecule data of cell samples encoded with DMO PSF. This open-source software can be accessed at https://github.com/Li-Lab-SUSTech/FD-DeepLoc. Furthermore, for in-situ single-molecule blinking data extraction and in-situ PSF modeling, as well as the computation of Zernike polynomial coefficients, we utilized our recently published open-source software, which can be accessed at https://github.com/ries-lab/uiPSF. We extracted single-molecule images from each optical section, which were acquired in a cyclic manner from the image stacks. These images were then merged and analyzed to generate a localization list for each section.

Despite having implemented an axial focusing system in the hardware setup (Fig. S1), the overall lateral drift of the sample and the accuracy of axial refocusing positions can result in slight misalignments of adjacent optical sections. This might lead to a deterioration of the resolution and distortions of the super-resolution images during the merging of sections. When performing drift correction, all sections were corrected using the drift values from the first section to prevent stitching artifacts and distortions in super-resolution images caused by computational offsets across different sections. The drift correction was performed using the redundant cross-correlation method[41] in SMAP[33]. When stitching data from neighboring optical sections, we employed the same redundant cross-correlation method as in drift correction. We scanned the entire cell sample in the axial direction with 1 µm step sizes, maintaining an optical section thickness of 1.2 µm. We ensured overlapping regions between adjacent optical sections to facilitate the use of the redundant cross-correlation method. Initially, we aligned neighboring optical sections in the XY direction, followed by alignment in the XZ direction. This approach enabled precise 3D alignment and stitching of the entire cell data.

### D. Biological sample preparation

#### 1. Cell culture

U2OS cells (Nup96-SNAP no. 300444, Cell Line Services) were grown in DMEM (catalog no. 10569, Gibco) containing 10% (v/v) fetal bovine serum (FBS; catalog no. 10099-141C, Gibco), 100 U/ml penicillin and 100 μg/ml streptomycin (PS; catalog no. 15140-122, Gibco) and 1× MEM NEAA (catalog no. 11140-050, Gibco). COS-7 cells (catalog no. 100040, BNCC) were grown in DMEM containing FBS and PS. Cells were cultured in a humidified atmosphere with 5% CO_2_ at 37 °C and passaged every two or three days. Before cell plating, high-precision 25-mm-round glass coverslips (catalog no. CG15XH, Thorlabs) were cleaned by sequentially sonicating in 1 M potassium hydroxide (KOH), Milli-Q water and ethanol, finally irradiated under ultraviolet light for 30 min. For super-resolution imaging, U2OS and COS-7 cells were cultured on the clean coverslips for 2 days with a confluency of ∼80%.

#### 2. SNAP-tag labeling of Nup96

To label Nup96, U2OS-Nup96-SNAP cells were prepared as previously reported[32]. Cells were prefixed in 2.4% (w/v) paraformaldehyde (PFA) for 30s, permeabilized in 0.4% (v/v) Triton X-100 for 3 min, and subsequently fixed in 2.4% PFA for 30 min. The buffer used for fixing and permeabilization was preheated to 37 °C before use. Then, cells were quenched in 0.1 M NH_4_Cl for 5 min and washed twice with PBS. To decrease unspecific binding, cells were blocked for 30 min with Image-iT FX Signal Enhancer (catalog no. I36933, Invitrogen). For labeling, cells were incubated in dye solution (1 μM SNAP-tag ligand BG-AF647 (catalog no. S9136S, New England Biolabs), 1 mM DTT (catalog no. 1111GR005, BioFroxx) and 0.5% (w/v) bovine serum albumin (BSA) in PBS) for 2 h and washed 3 times in PBS for 5 min each to remove excess dyes. Lastly, cells were postfixed with 4% PFA for 10 min, washed with PBS 3 times for 3 minutes each, and stored at 4 °C until imaged.

#### 3. Microtubule labeling

Microtubule samples were prepared as previously described[30]. COS-7 cells were prefixed with 0.3% (v/v) glutaraldehyde (GA) and 0.25% Triton X-100 in the cytoskeleton buffer [CB (10 mM MES, 5 mM glucose, 150 mM NaCl, 5 mM MgCl_2_, 5 mM EGTA, pH 6.1), preheated to 37 °C before using)] for 1-2 min. Removed waste liquor and fixed with 2% GA in CB for 10min (preheat to 37 °C before using). Remove waste liquor and quenched in 0.1% (w/v) NaBH_4_ (0.01g in 10 ml PBS) for 7 min and washed 3 times for 5 minutes each with PBS. Then incubated in the permeabilization buffer (0.1% Triton X-100 in 3% BSA) for 10min. After washing 3 times for 5 minutes each with PBS, cells were stained by goat anti-mouse β-tubulin (catalog no. T4026, Sigma, 2.4 mg/ml) with 1:1,000 dilution in 3% BSA for 1 h and washed 3 times for 5 minutes each with PBS. Cells were then stained with the corresponding secondary antibodies conjugated with AF647 (catalog no. 2289596, Invitrogen, 2 mg/ml) with 1:1,000 dilution in 3% BSA for 1 h and washed with PBS. Finally, cells were postfixed with 4% PFA for 10 min, washed with PBS 3 times for 5 minutes each, and stored in PBS at 4 °C.

#### 4. Mitochondria labeling

Mitochondrial samples were prepared as previously described[42]. COS-7 cells were fixed with 4% PFA (preheat to 37 °C before using) in PBS for 12 min, incubated in permeabilization buffer (0.3% CA-630 (catalog no. I8896, Sigma), 0.05% TX-100, 0.1% BSA and 1× PBS) for 3 min, and then quenched in 0.1 M NH_4_Cl for 5 min. After washed 3 times for 5 min each with PBS, cells were blocked in 3% BSA for 60 min. For labeling, cells were stained by rabbit anti-Tom20 (catalog no. ab78547, Abcam, 1 mg/ ml) with 1:1,000 dilution in 3% BSA for 2 h and washed 3 times for 5 min each with PBS. Cells were then stained with the corresponding secondary antibodies conjugated with AF647 (catalog no. A21245, Invitrogen, 2 mg /ml) with 1:2,000 dilution in 3% BSA for 2 h, and washed 3 times for 5 min each with PBS. Finally, cells were post-fixed with 4% PFA for 10 min, washed with PBS 3 times for 5 min each, and stored in PBS at 4 °C.

#### 5. Imaging Buffer

Samples were imaged in refractive index matching buffer, including 50 mM Tris-HCl (pH 8.0), 10 mM NaCl, 10% (w/v) glucose, 0.5 mg/ml glucose oxidase (G7141, Sigma), 40 μg/ml catalase (C100, Sigma), 35 mM cysteamine, and 28.5% (v/v) 2,2’-thiodiethanol (166782, Sigma). The refractive index of the final imaging buffer is 1.406.

#### 6. Beads preparation

We first diluted the 100-nm-diameter crimson beads (custom-designed, Invitrogen) to 1:40,000 in Milli-Q water and vortexed the mixture for 3-5 minutes. Then, we pipetted 40 μl of 1 M MgCl_2_ in the center of the 25-mm-diameter coverslip (cleaning protocol refers to the cell culture section) and mixed with 360 μl of the diluted bead solution on the coverslip. The mixture was incubated for 5 min, washed 3 times with Milli-Q water, and stored in Milli-Q water at 4 °C in the dark.

## Acknowledgments

The authors thank Shuang Fu and Mengfan Li for helpful suggestions on DM calibration and PSF optimization.

## Data availability

Data are available upon reasonable request.

## Supplemental document

See Supplement 1 for supporting content.

