## supplemental1 for "Aberration correction for deformable mirror based remote focusing enables high-accuracy whole-cell super-resolution imaging"

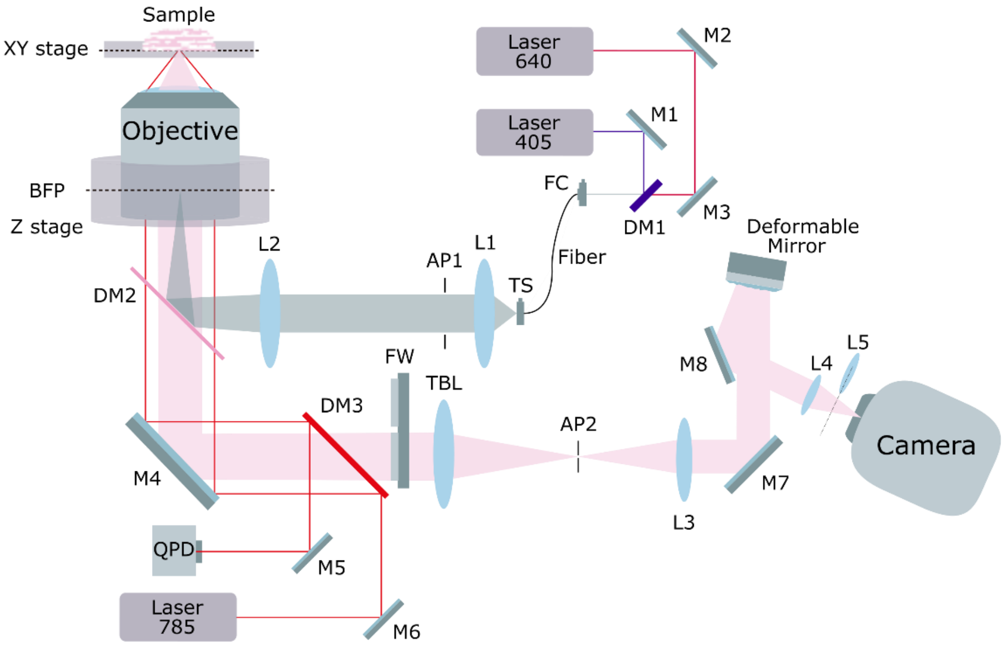


**Fig. S1.** Detailed layout of the optical setup. M: mirror, DM: dichroic mirror, L: lens, TS: translation stage, FC: fiber coupler, Fiber: single-mode fiber, BFP: back focal plane, FW: filter-wheel, TBL: tube lens, AP: aperture, QPD: quadrant photodiode. The excitation lasers are first reflected by dichroic mirror DM1, then coupled into a single-mode fiber through the fiber coupler FC. Before being reflected by the main dichroic mirror DM2 to enter the objective for sample illumination, a pair of lenses L1 and L2 along with a slit at AP1 are used for collimating and reshaping the beam. In the imaging path, the fluorescence collected by the objective is transmitted through the dichroic mirror DM2 and filtered by the filters on the filter wheel FW. It is then imaged onto the aperture AP2, which is confined by a slit, using a tube lens TBL. Subsequently, the fluorescence passes through a 4f system composed of lenses L3 and L4, with a deformable mirror installed in the Fourier plane, and ultimately, the fluorescence signal is detected by the camera. Additionally, a beam excited by a 785 nm laser, reflected off the coverslip, is detected by the quadrant photodiode QPD, which provides feedback control to the Z stage for focus locking.

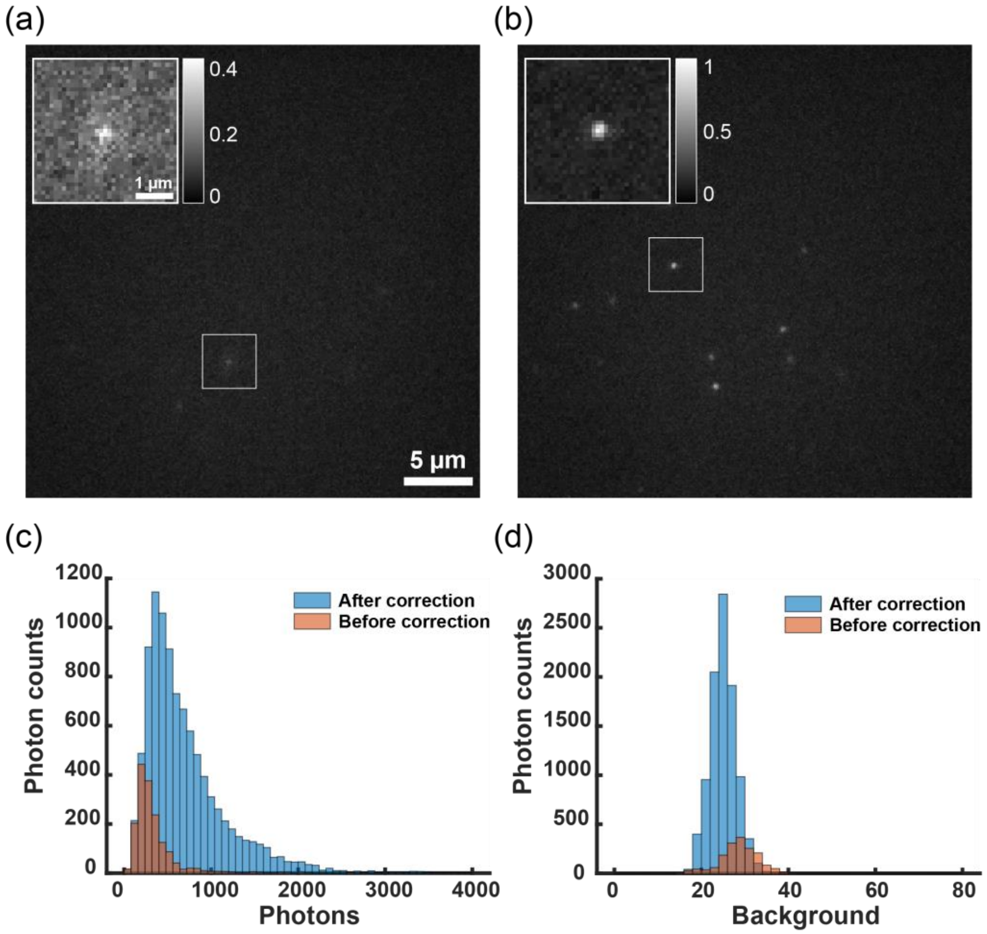


**Fig. S2.** Comparison of raw single-molecule images before and after aberration correction, accompanied by the distribution of photons and background photons. (a) Snapshots of an unmodified 2D single-molecule image of Nup96-SNAP-AF647 labeled NPCs in U2OS cell samples refocused to 4 μm above the nominal focal plane. The inset shows a zoomed view of the fluorophore in the boxed area. (b) Snapshots of the 2D single-molecule image after aberration correction, with a significantly improved SNR in the image. The inset clearly shows the successful recovery of the distorted PSF. (c) The photon distribution analysis from 2,000 frames of 2D single-molecule images before and after aberration correction reveals that 1,644 single-molecule events with an average photon count of 335 were localized before correction, whereas 9,680 single-molecule events with an average photon count of 711 were localized after correction. (d) The background photon distribution analysis from the same 2,000 frames of 2D single-molecule images before and after aberration correction showed an average background photon count of 29 before correction and an average photon count of 25 after correction. The single-molecule raw images were acquired at 120 mW laser power and 20 ms exposure time.

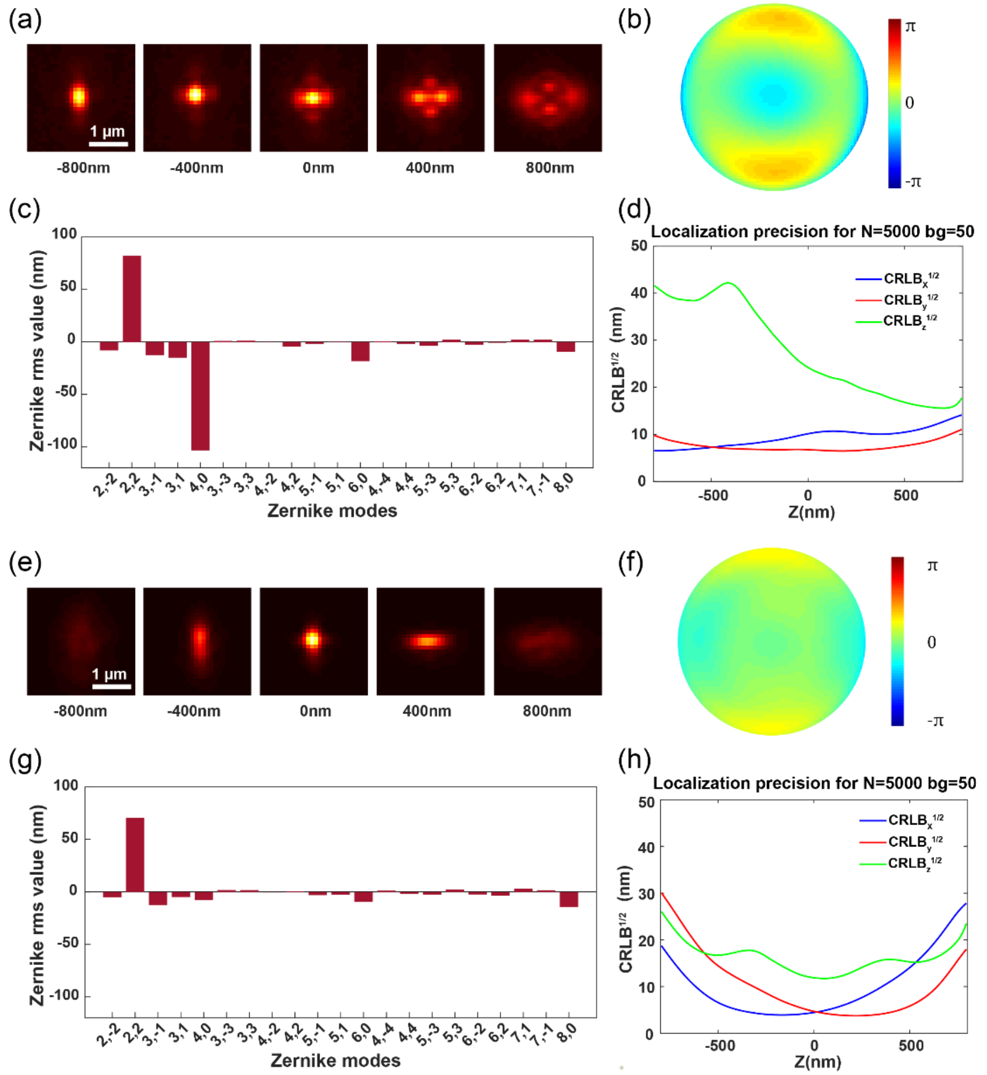


**Fig. S3.** Correcting aberrations during refocusing with astigmatism PSF using a DM at various axial positions of the beads on the coverslip. (a) The beads are refocused using a DM to a position 2 μm above the nominal focal plane. (b) The pupil function for the refocused beads. (c) Fitted 21 Zernike coefficients for the beads stack. (d) CRLB for the experimental astigmatism PSF, with 5,000 photons and 50 background photons used in the CRLB calculation. (e-h) The PSF shape, pupil function, Zernike coefficients, and localization precision after aberration correction, respectively. The CRLB exhibits significant improvements, particularly in the Z-axis localization precision.

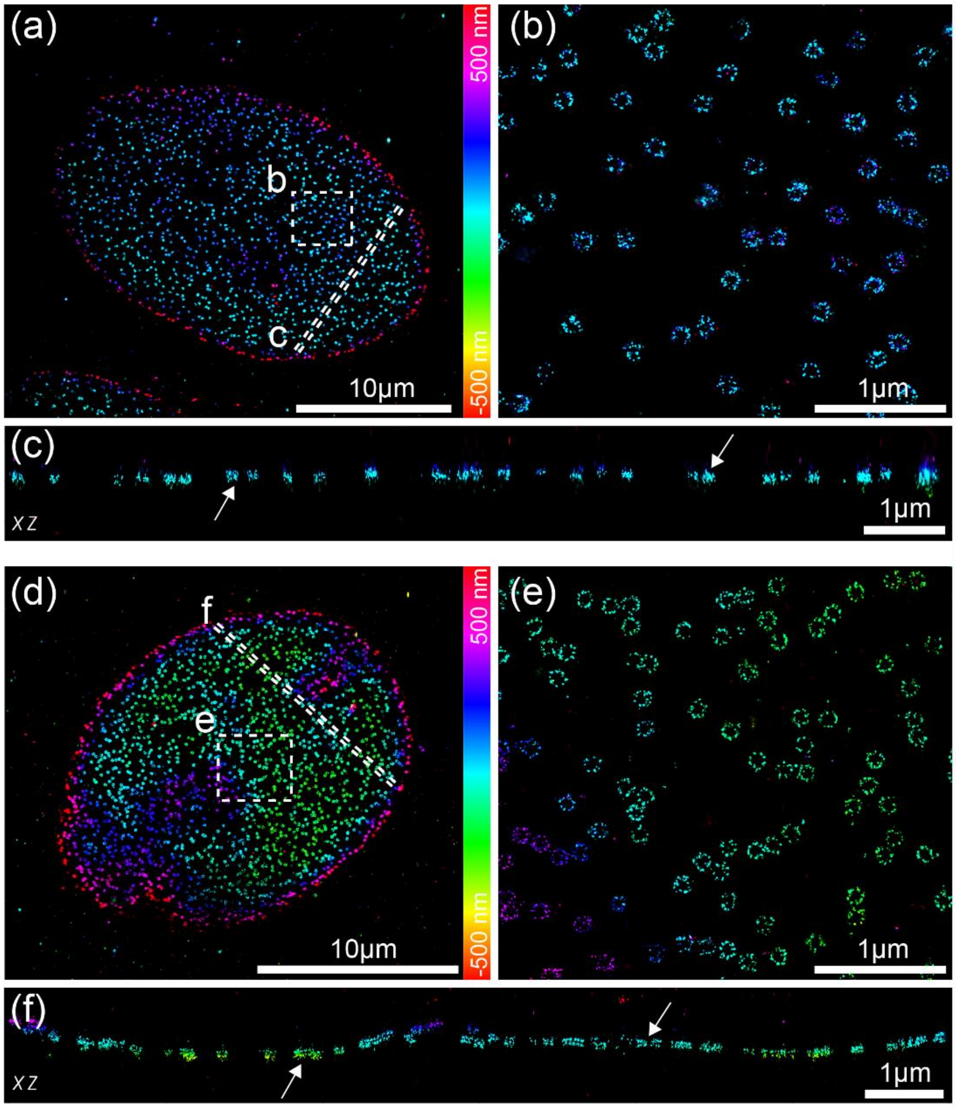


**Fig. S4.** Experimental astigmatic 3D data of NPCs before and after aberration correction by refocusing. (a) Overview of 3D super-resolution image of Nup96-SNAP-AF647, refocused using a DM at 2 μm above the nominal focal plane. Reconstructed using the Cspline experimental PSF model shown in Fig. S3(a). (b) Magnified view of the area denoted by the box in a. (c) Side-view cross-section of the region denoted by the dashed line in a. (d-f) 3D super-resolution image of NPCs after aberration correction at the refocused position. Significant improvements were observed in the reconstructed images, particularly in the XZ plane, where the double-ring structure of NPCs became distinctly visible after aberration correction, as indicated by the white arrows in panels c and f. The data were acquired from 50,000 frames with 120 mW laser power and 20 ms exposure time.

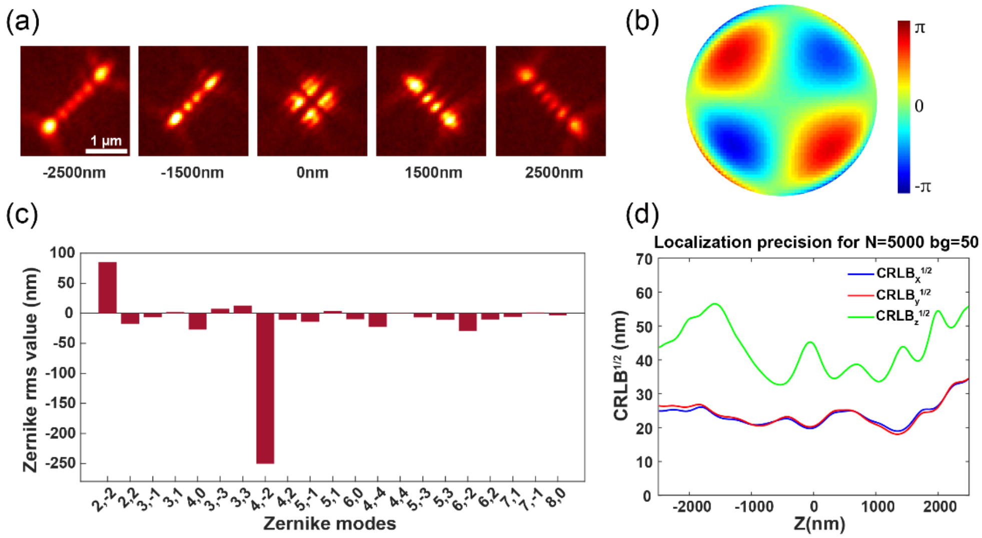


**Fig. S5.** Optimization of tetrapod PSF using a DM for the beads on the coverslip. (a) The averaged tetrapod PSF for fluorescent beads on the coverslip, with PSFs at axial positions from -2500 nm to 2500 nm. (b) The pupil function for the beads. (c) Fitted 21 Zernike coefficients for the beads stack. (d) CRLB for the experimental tetrapod PSF, with 5,000 photons and 50 background photons used in the CRLB calculation.

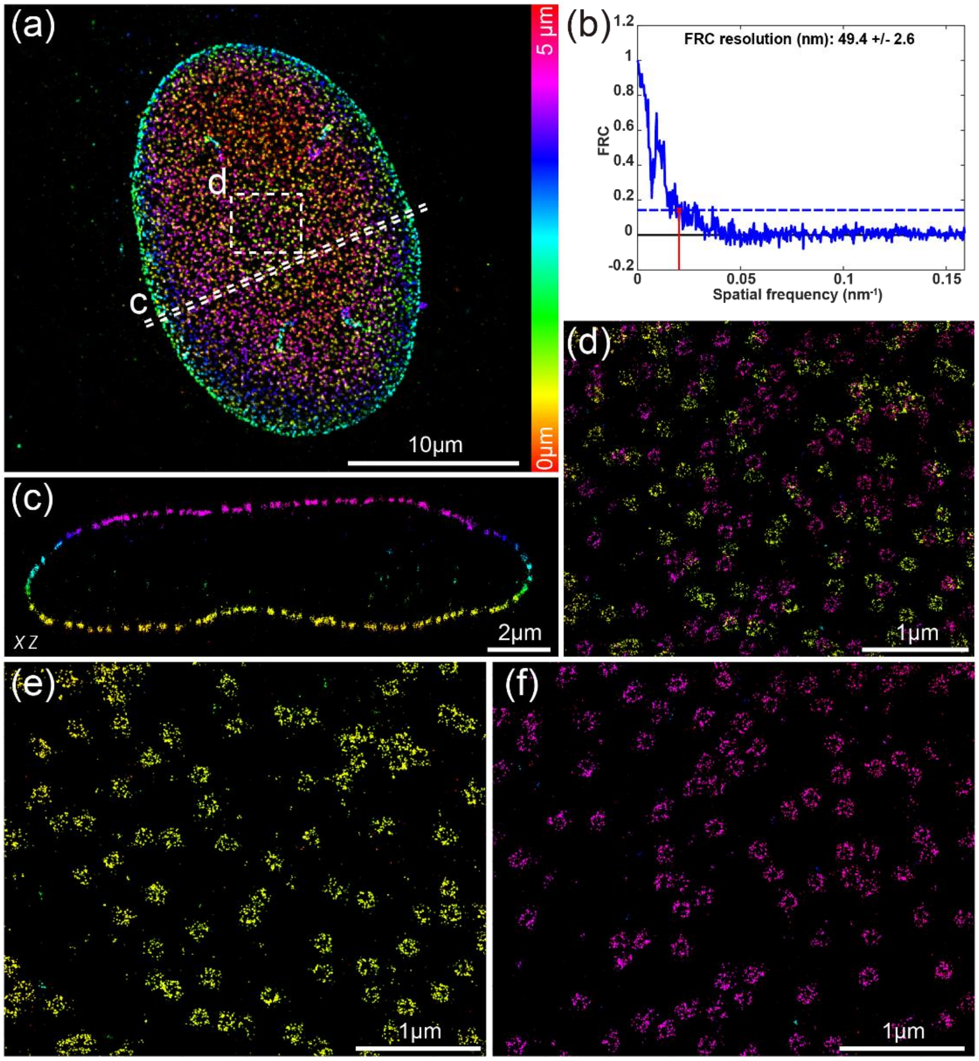


**Fig. S6.** Whole-cell 3D super-resolution imaging of NPC using DMO tetrapod PSF. (a) Overview of the panoramic whole-cell 3D imaging of NPC using DMO tetrapod PSF without remote focusing. (b) FRC analysis of the regions indicated in a. (c) Side-view cross-section of the region denoted by the dashed line in a. (d) Magnified view of the area denoted by the box in a. (e) View of the bottom surface of the area in d. (f) View of the top surface of the area in d. The data were acquired from 100,000 frames with 120 mW laser power and 20 ms exposure time.

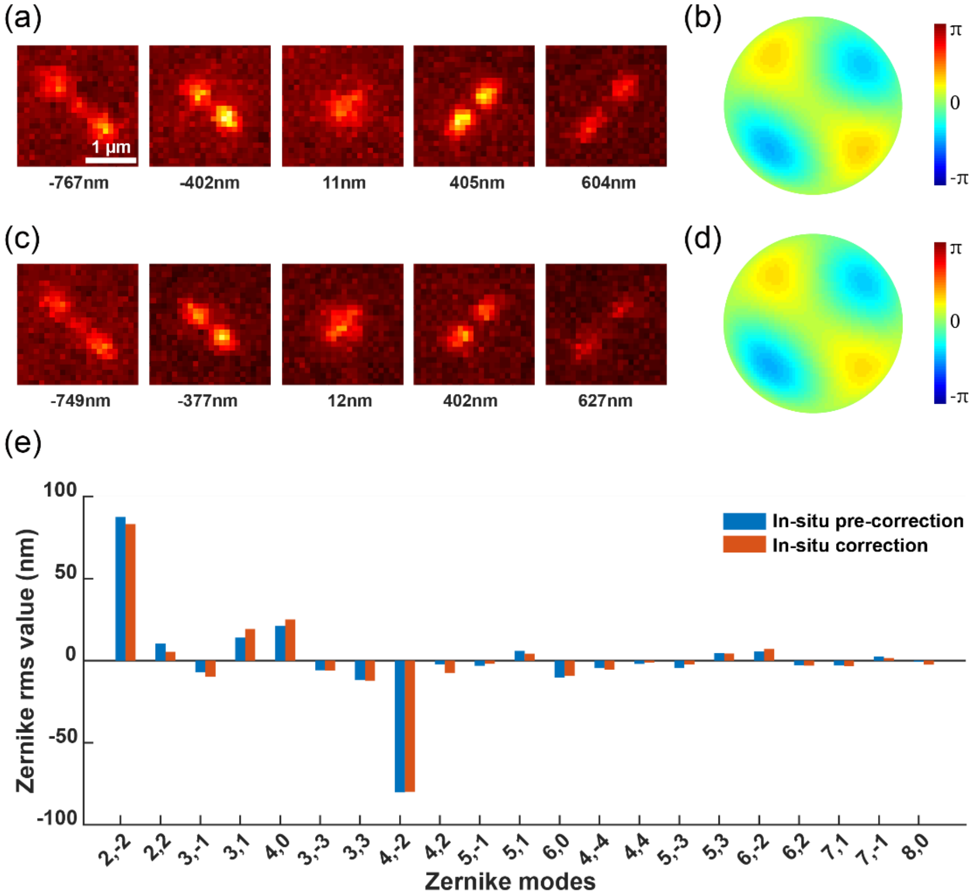


**Fig. S7.** Comparison of in-situ aberration pre-correction and actual aberration correction differences in various cells using single-molecule blinking data. (a) Pre-correction PSF derived from single-molecule blinking data of immunofluorescence-labeled TOM20 in COS7 cells at a depth of 2.5 μm. (b) Pupil function calculated from the in-situ PSF of pre-correction. (c-d) The PSF shape and its corresponding pupil function after in-situ aberration correction. (e-f) The fitted 21 Zernike coefficients obtained from the in-situ single-molecule blinking data of pre-corrected cells, contrasting the differences between pre-corrected and actual corrected cells.
